# Contextual interference effect is independent of retroactive inhibition but variable practice is not always beneficial

**DOI:** 10.1101/466896

**Authors:** Benjamin Thürer, Sarah Gedemer, Anne Focke, Thorsten Stein

**Affiliations:** Brain Signaling Group, Institute of Basic Medical Sciences, Department of Molecular Medicine, University of Oslo, Oslo, Norway; BioMotion Center, Institute of Sports and Sports Science, Karlsruhe Institute of Technology, Karlsruhe, Germany

**Author notes:** Correspondence: Dr. Benjamin Thürer.

**Keywords:** Motor memory consolidation, Force field adaptation, Sensorimotor learning, Motor adaptation, Retrograde inhibition, Contextual interference, Variable practice

## Abstract

Positive effects of variable practice conditions on subsequent motor memory consolidation and generalization are widely accepted and described as the contextual interference effect (CIE). However, the general benefits of CIE are low and these benefits might even depend on decreased retest performances in the blocked-practicing control group, caused by retroactive inhibition. The aim of this study was to investigate if CIE represents a true learning phenomenon or possibly reflects confounding effects of retroactive inhibition. We tested 48 healthy human participants adapting their reaching movements to three different force field magnitudes. Subjects practiced the force fields in either a Blocked (B), Random (R), or Constant (C) schedule. In addition, subjects of the Blocked group performed either a retest schedule that did (Blocked-Matched; BM) or did not (Blocked-Unmatched; BU) control for retroactive inhibition. Results showed that retroactive inhibition did not affect the results of the BU group much and that the Random group showed a better consolidation performance compared to both Blocked groups. However, compared to the Constant group, the Random group showed only slight benefits in its memory consolidation of the mean performance across all force field magnitudes and no benefits in absolute performance values. This indicates that CIE reflects a true motor learning phenomenon, which is independent of retroactive inhibition. However, random practice is not always beneficial over constant practice.

## 1. Introduction

It is widely accepted that variable practice conditions can be beneficial for motor memory consolidation (Schmidt, 1975; Shea & Morgan, 1979). In particular, the contextual interference effect (CIE) – originally formulated by Battig (1972) for verbal learning – describes an increased retest and transfer performance (Shea & Morgan, 1979; Magill & Hall, 1990) due to highly-interfering cognitive processes during random practice conditions (Kantak et al., 2010; Lage et al., 2015). This effect seems to be more robust in basic research than in applied settings (Brady 2004, Brady 2008). In addition, CIE is commonly examined by comparing a random with a blocked practice schedule and test for their corresponding effects on posttest and transfer test performance. In such a random practice schedule, different tasks (or parameters) change randomly from trial to trial, whereas, in a blocked practice schedule, one specific task (or parameter constellation) is practiced as a whole block first, before switching to the next task (or parameter constellation).

Although CIE seems to be robust, there is no widely accepted hypothesis that accounts for this effect. Classical explanations include the elaboration hypothesis (Shea & Zimny, 1983), the reconstruction hypothesis (Lee & Magill, 1983), and the retroactive inhibition hypothesis (Shea & Titzer, 1993). Thus, it is still unsolved whether CIE stems from an increased memory consolidation due to the random practice condition (e.g. elaboration or reconstruction hypothesis) or by a decreased retention performance of the blocked practice condition (retroactive inhibition hypothesis). This latter assumption – which is in the focus of this paper – derives from the observation that subsequent learning of different tasks can lead to inhibition of a previous memory, an effect called retroactive inhibition (see Robertson et al., 2004 for a review). Concerning CIE, retroactive inhibition might lead to disadvantages for the blocked practicing subjects since their previous memory might be inhibited due to the blocked practice schedule. Therefore, these subjects might show the worst performance when recalling the first task and the best performance when recalling the last task they have practiced. Previous work showed possible confounding effects of this retroactive inhibition on the motor retrieval after blocked practice and, therefore, questioned the validity of CIE (Del Rey et al., 1994, Shewokis et al., 1998). Furthermore, when retroactive inhibition was eliminated by using a reminder trial, no differences in memory recall were observed between random and blocked practice schedules (Shea & Titzer, 1993).

So far, CIE and retroactive inhibition were discussed in the context of skill learning, in which most CIE studies were conducted. Skill learning is commonly defined as a “set of processes associated with practice or experience leading to relatively permanent changes in the capacity for skilled movement” (Schmidt et al., 2018, p. 283). In contrast, motor adaptation is interpreted as a different type of motor learning, in which the motor system responds to changes in environmental conditions and/or changes in the body to regain the former capacity for a skilled movement under these new conditions (Krakauer & Manzoni, 2011). This study focuses on motor adaptation using a force field paradigm (Shadmehr & Mussa-Ivaldi, 1994), for which CIE has been demonstrated in previous studies from our laboratory (Thürer et al., 2017; Thürer & Weber et al., 2018). In these studies, subjects had to adapt their reaching movements to different force field magnitudes either in a blocked or random fashion. However, these former studies did not control for retroactive inhibition and a constant group, practicing only the force field magnitude that needed to be recalled, was not included.

Therefore, the first purpose of this study is to control for the confounding effects of retroactive inhibition and examine the validity of the contextual interference effect in force field adaptation. The second purpose of this study is to examine if variable practice schedules (blocked and random) outperform a constant practice schedule even if subjects of the constant group have the advantage of adapting their reaching movements only to a single force field.

## 2. Methods and Methods

### 2.1. Participants

This study tested 48 healthy right-handed participants (24 ± 4 years; 10 women) with no previous experience at a robotic manipulandum. Handedness was tested by the Edinburgh inventory (Oldfield, 1971) and participant’s vision was normal or corrected to normal. The study was approved by the Institutional Review Board. All participants were informed about the protocol and gave their written informed consent in accordance with the Declaration of Helsinki.

### 2.2. Apparatus and experimental task

The experimental task was implemented at a robotic manipulandum (Kinarm End-Point Lab, BKIN Technologies, Kingston, Canada) which can produce forces via a handle towards the participants’ hands. In addition, we used a virtual reality display that allowed the participants to see the visual information in the horizontal plane (Fig 1A). Please note that vision of handle, hand, and arm was occluded by the virtual reality display. Positions and forces of the robot handle were sampled at 1000 Hz.

**Figure 1:**
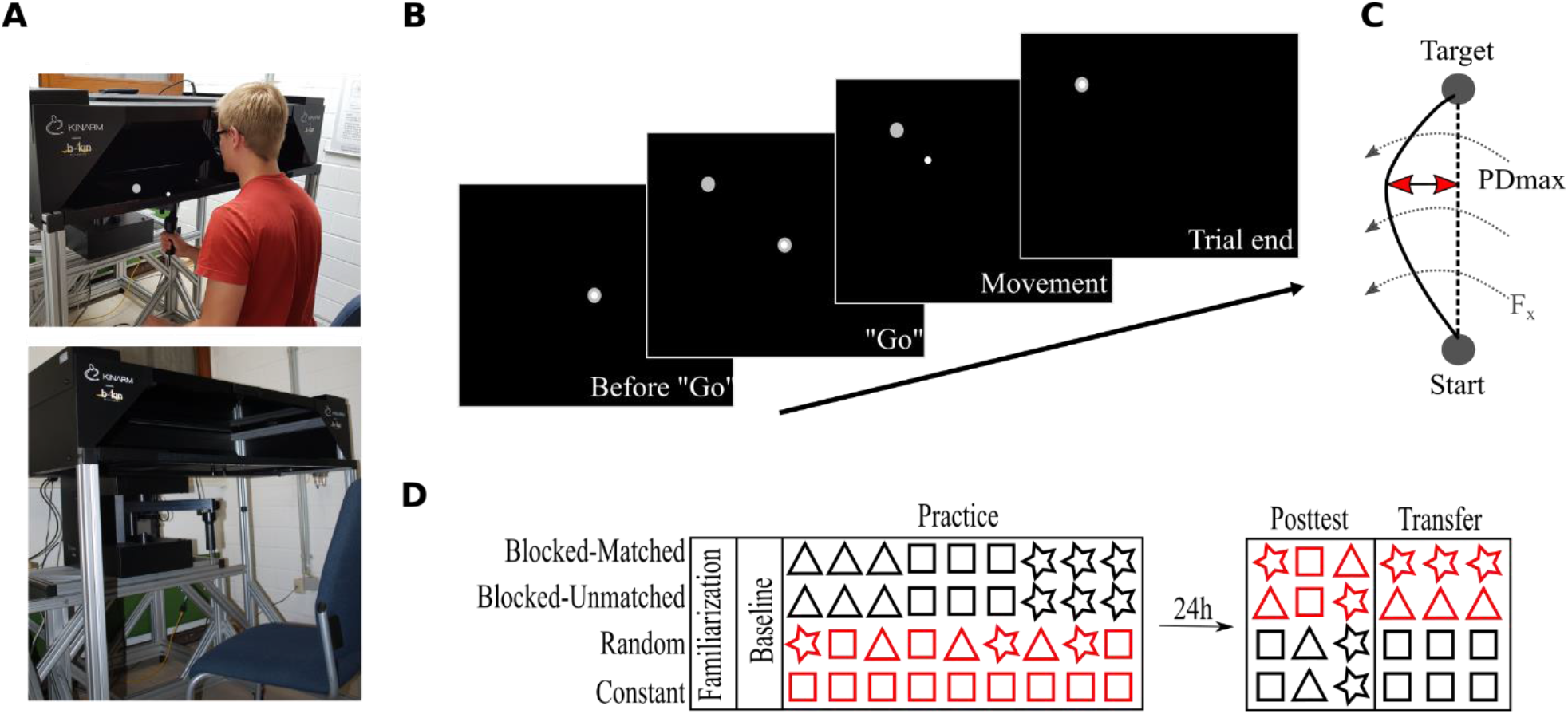
Apparatus and task. **A**: Robotic manipulandum with the virtual reality display. Subjects hold the robotic handle but look into the virtual reality add-on in front of them. In addition, a fabric, not shown here, was attached to the reality add-on and to the subjects’ shoulders to prevented further visual input of the arms. The virtual cursor is controlled via the robotic handle. Permission to publish this figure was given by the pictured person. **B**: Experimental task at the robotic manipulandum. The subjects see the cursor and targets on the screen in the horizontal plane. **C**: the maximum perpendicular displacement (PDmax) was used to quantify the motor performance. Dashed arrows indicate the force field. **D**: Experimental protocol over two consecutive days. Red colors indicate that both Blocked groups differed in their retest schedule on day 2 whereas the Random and Constant groups differed in their practice schedules on day 1. Triangles, rectangles, and stars symbolize different force field magnitudes.

We will briefly describe the experimental task which can be found elsewhere in more detail (Thürer et al., 2017). Participants were seated in front of the manipulandum and the virtual reality display was calibrated to the robot’s handle. All participants performed center-out reaching movements with their dominant (right) hand. While performing this task, the horizontal display shows a white cursor which is controlled via the handling of the manipulandum (Fig. 1B). Every trial started by holding the cursor in the center target on the screen and a “go” signal was given by the highlighting of a target. From that “go” signal on, participants were allowed to start their reaching movement without any pressure of time (no fast reaction times required). When participants reached the target position, subjects were actively moved backwards to the center position by the manipulandum. After a short pause in the center position of 800 ms, the next target highlighted in a pseudo-randomized order. In total, six target positions were defined building a circle with a diameter of 20 cm surrounding the center point. Pseudo-randomization facilitated that in every block of six trials every target highlighted just once and that every participant had a different target order so that no influence of target direction was given on the group level.

To provide similar movement times across trials and subjects, visual feedback was implemented in every single trial. The feedback was given via a change in the target color after reaching it. Target color switched to red if the movement was too fast (< 450 ms), blue if it was too slow (> 550 ms), and green otherwise.

To induce motor adaptation and subsequent memory consolidation, we implemented velocity-dependent counter-clockwise directed force fields at the robotic manipulandum. These force fields perturbed the participants’ movements and typically degraded their initial motor performance leading to curved hand trajectories (Fig. 1C). In order to investigate practice schedules with different amounts of variabilities, three separate force field viscosities were implemented with each viscosity inducing a force field magnitude of either 8, 15, or 22 Ns/m. Therefore, each force field magnitude represented an object with different physical properties. The absolute maximum perpendicular displacement between the participant’s hand path and a direct line joining center point and target quantified the motor error (Fig. 1C).

### 2.3. Experimental procedure

Participants were equally distributed into 4 groups (Blocked-Matched, BM; Blocked-Unmatched, BU; Constant, C; Random, R; each *n* = 12). The groups differed only in their task protocol during Practice and during Retest (Posttest and Transfer). The study took place on two consecutive days with 24 h between the two test sessions (Fig. 1D).

On day 1, all participants received instructions about the behavioral task and performed 144 familiarization trials under null field conditions (motors of the robot were turned off) with two breaks of 30 s after every 48th trials. Then, participants performed a baseline measurement consisting 30 null field trials. After that, all participants performed 540 force field trials during Practice, with a different force field schedule according to their group allocation. To avoid fatigue, participants had a 30 s break after each 60th trial. The participants performed all trials on day 1 with their dominant right hand.

The practice schedule was identical between the two Blocked groups (BM, BU) but different for the Random and Constant groups. Participants of the Blocked groups performed the three force field magnitudes (8, 15, 22 Ns/m) in a blocked order. Therefore, all trials of one specific magnitude were practiced first, before switching to the next magnitude. This resulted in three blocks, each containing 180 trials of one specific force field magnitude. The Random group performed a highly-variable practice schedule so that the three force field magnitudes changed on a single-trial level. For the Constant group, each participant practiced only one specific force field magnitude (e.g. 15 Ns/m) and, thus, encountered no force field variability at all. The force field magnitude (for C) and the magnitude order (for BM, BU, and R) was counter-balanced across participants so that the mean force field magnitude was 15 Ns/m on the group level. In addition, for the Blocked and Random groups, the mean force field magnitude across the whole Practice session was 15 Ns/m for each single participant.

On day 2, all participants performed a Posttest and Transfer test. To quantify Posttest performance, all participants performed 18 force field trials divided into three blocks with each block representing one force field magnitude. Then, participants performed 60 trials of a constant force field magnitude with their non-dominant left hand (Transfer test) to investigate long-term effects on the contralateral hand indicated by a previous study from our group (Thürer & Weber et al., 2018).

The order of force field magnitudes on day 2 differed between groups. For the Blocked-Matched group, the magnitudes in Posttest were in a reversed order compared to Practice and, thus, “matched” in terms of a reduced effect of retroactive inhibition on the first block of the retest schedule (Fig. 1D). For instance, when for a specific participant the Practice order was 15, 8, 22 Ns/m, the Posttest order was set to 22, 8, 15 Ns/m. For the Blocked-Unmatched group, however, the order of force field magnitudes was the same for the Practice and the Posttest session. For instance, when for a specific participant the Practice order was 15, 8, 22 Ns/m, the Posttest order was set to 15, 8, 22 Ns/m. The order of magnitudes on day 2 was similar for the Random and for the Constant group. Both groups started the Posttest with the force field magnitude (for C) or with the mean force field magnitude (for R, i.e. 15 Ns/m) of the Practice session. This is due to a study that has shown that participants adapt to the mean force field magnitude (Scheidt et al., 2001). Regarding the Constant group, the first block’s magnitude was different between participants for each single participant had a different magnitude during Practice due to counterbalancing. Both groups (R, C) were counterbalanced for the order of the remaining two force field magnitudes so that, still, the mean across groups for the second and third block of the Posttest was at 15 Ns/m. According to the Posttest, all participants performed the Transfer test on the left hand at a specific constant force field magnitude, which was the same as the first magnitude in the Posttest (Fig. 1D).

### 2.4. Statistics

For the statistical analyses, mean performance for the first and the last 6 trials of the Practice session (Practice FT, Practice LT) was computed. Posttest performance was computed by the mean of the first, middle, and last 6 Posttest trials (Posttest FT, MT, LT) and the mean across all 18 Posttest trials (Posttest ALL). Contralateral Transfer performance was quantified by the initial 6 Transfer trials (Transfer FT) and the whole 60 Transfer trials (Transfer ALL).

To test for the possible influence of retroactive inhibition on CIE, we performed mixed-model 2*2 ANOVAs with the factors time (Practice LT, Posttest FT; Practice LT, Posttest ALL) and group (BM, BU). For a possible effect on the generalization from one hand to the other, the factor time was adjusted accordingly (Practice LT, Transfer FT; Practice LT, Transfer ALL). In addition, we investigated if random practice even outperforms constant practice by using standard Fischer t-tests between groups (R, C). Therefore, we calculated differences between each force field magnitude of the Posttest (Posttest FT, MT, LT, ALL) and the last trials of the Practice session (Practice LT), respectively.

It is widely accepted that p-values alone are not a good marker for potential results in research (e.g. Nuzzo, 2014). Therefore, besides using classical inferential statistics, we provide effect sizes (partial eta squared, *pEta^2^*; Cohen’s d, *d*) and additional Bayesian statistics. Bayesian statistics are provided in the supplementary material and confirm the results and interpretations of the classical inferential statistics.

All parameters were tested for normal distribution and homogeneity of variances using Shapiro-Wilk and Levene test. Statistical analyses were performed using MATLAB R2015b (Mathworks, Natick, USA) and JASP 0.8.6 (Team JASP). Threshold for statistical significance was set to *p* = 0.05. Multiple comparisons were corrected by the False Discovery Rate (FDR, Benjamini & Hochberg, 1995) and in case of multiple comparisons, p-values in this study represent the FDR corrected p-value (Benjamini & Yekutieli, 2001).

## 3. Results

### 3.1. CIE is unaffected by the different retest schedules of the Blocked groups

The progress in motor performance for both Blocked groups is depicted in Fig. 2A. First, we tested if motor adaptation during Practice differed between groups. Both groups (BM, BU) adapted successfully to the force field schedule during Practice (*F*(1,22) = 104.09, *p* < .001, *pEta^2^* = 0.83, for the factor time (Practice FT, Practice LT)) and showed no differences in their adaptation (*F*(1,22) = 0.79, *p* = .385, *pEta^2^* = 0.03 for the factor group (BM, BU); *F*(1,22) = 1.67, *p* = .210, *pEta^2^* = 0.07, for mixed-model ANOVA with time*group interaction).

**Figure 2:**
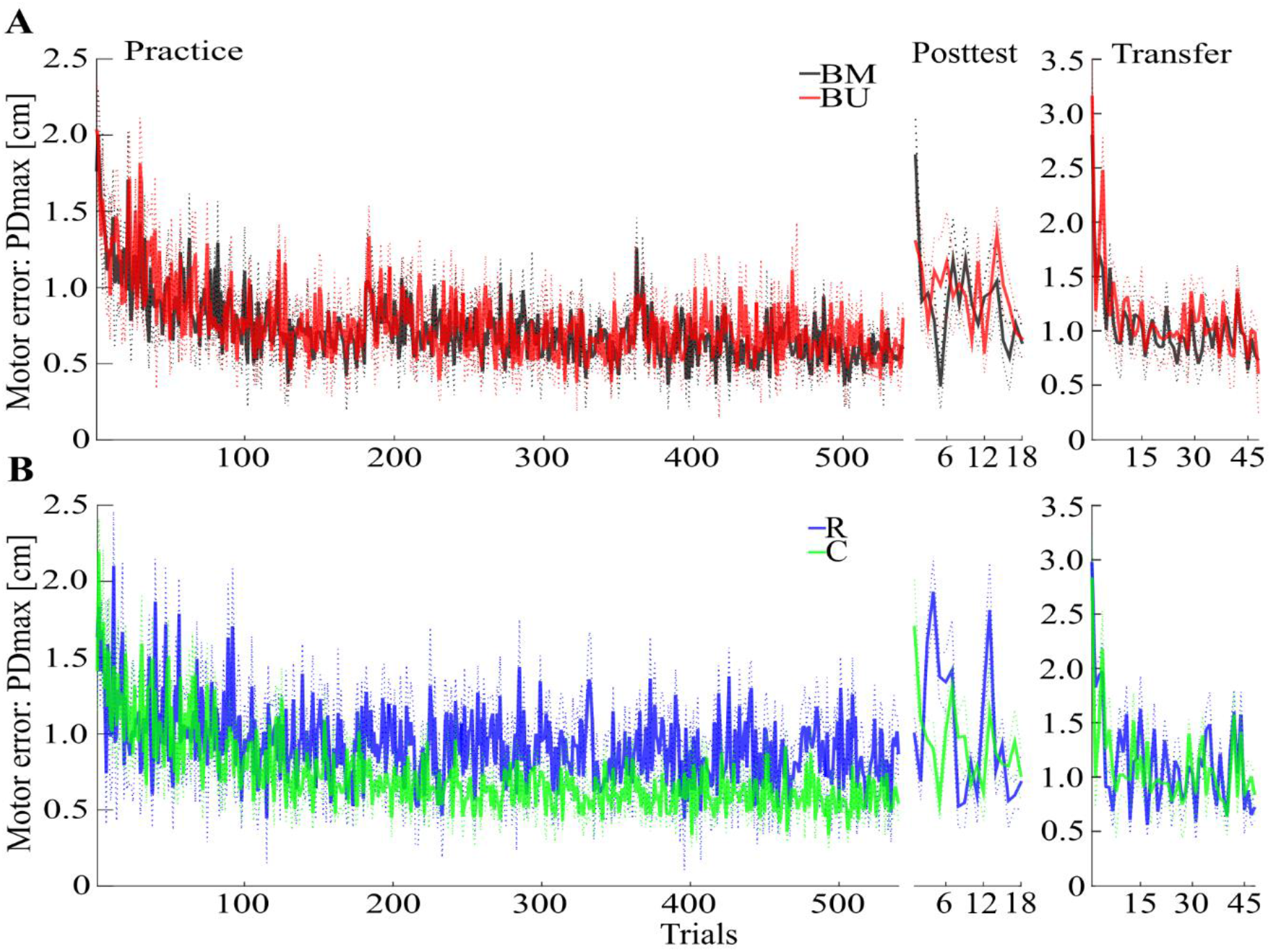
Descriptive results. **A**: Progress of the mean motor error with SEM for Practice, Posttest, and Transfer test of the Blocked-Matched (BM) and the Blocked-Unmatched (BU) group. **B**: Progress of the mean motor error with SEM for the Random (R) and the Constant (C) group.

Consolidation of motor memory (from Practice to Posttest) did not differ between Blocked groups regarding their recall of the first force field magnitude (*F*(1,22) = 0.47, *p* = .498, *pEta^2^* = 0.02) or regarding all force field magnitudes (*F*(1,22) = 0.06, *p* = .808, *pEta^2^* < 0.01, for uncorrected time*group interactions with factors time (Practice LT, Posttest FT; Practice LT, Posttest ALL) and group (BM, BU)). Although descriptive statistics indicate slight benefits for the BM group in recalling the very first force field magnitude during Posttest (Fig 2A, Fig 3A), this is not supported by additional post-hoc statistics (*t*(22) = −0.94, *p* = .358, *d* = −0.38, for uncorrected independent t-test between groups’ Posttest FT performance). However, memory consolidation was significantly stronger for the Random group compared to the BM group (*F*(1,22) = 5.65, *p* = .029, *pEta^2^* = 0.20) and descriptively stronger to the BU group (*F*(1,22) = 5.49, *p* = .054, *pEta^2^* = 0.20, for FDR corrected time*group interactions with factors time (Practice LT, Posttest ALL) and group (BM, R; BU, R)), which confirms the contextual interference effect.

**Figure 3:**
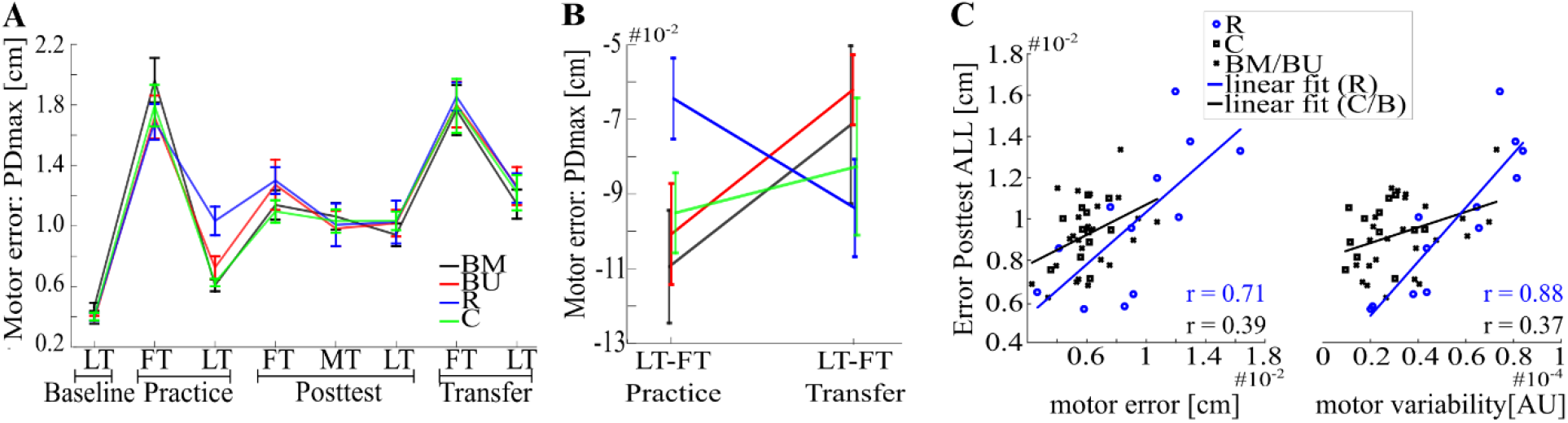
Deeper investigation of the behavioral results. **A**: Overview of the mean motor error with SEM computed across trials and subjects for each group over the whole experiment. **B**: Differences for Practice LT - Practice FT and Transfer LT - Transfer FT with SEM as an indication for the range of motor adaptation. **C**: Correlation analysis for motor error of Practice LT (left) and motor variability of Practice LT (right), each associated with Posttest ALL. Lines indicate linear fits for the Random (blue) and all the other groups (black).

We further investigated if retroactive inhibition affected the generalization from the dominant (Practice) to the non-dominant (Transfer) hand. No differences between Blocked groups (*F*(1,22) = 0.15, *p* = .701, *pEta^2^* < 0.01, for time*group interaction with factors time (Practice LT, Transfer FT) and group (BM, BU)) were observed and the Random group performed similar to both Blocked groups (BM vs. R: *F*(1,22) = 1.18, *p* = .568, *pEta^2^* = 0.05; BU vs. R: *F*(1,22) = 0.34, *p* = .580, *pEta^2^* = 0.02 for FDR corrected time*group interactions with factors time (Practice LT, Transfer FT) and group (BM, R; BU, R)). However, further adaptation (quantified by Transfer LT - Transfer FT) was observed to be faster for the Random group (Fig 3B), with this effect becoming more pronounced if adaptation during Practice (Practice LT - Practice FT) is compared with adaptation during Transfer (BM vs. R: *F*(1,22) = 3.39, *p* = .079, *pEta^2^* = 0.13; BU vs. R: *F*(1,22) = 8.05, *p* = .020, *pEta^2^* = 0.27, for FDR corrected time*group interaction with factors time (Practice LT – Practice FT, Transfer LT – Transfer FT) and group (BM, R; BU, R)). This indicates that, although motor memory generalization from Practice to Transfer did not differ between groups, adaptation within the Transfer test appears to be faster for the Random compared to both Blocked groups.

### 3.2. Random practice improves mean memory consolidation of multiple force field magnitudes

The second aim of this study was to examine if constant practice leads to better memory consolidation of only one force field magnitude than random practice and if random practice outperforms constant practice in the recall of multiple force field magnitudes. Our results show that memory consolidation of the Constant and of the Random group did not differ for each single force field magnitude (quantified by differences between Posttest FT/MT/LT and Practice LT. Posttest FT: *t*(22) = 0.89, *p* = .386, *d* = 0.36; Posttest MT: *U* = 99, *p* = .171; Posttest LT *t*(22) = 1.97, *p* = .122, *d* = 0.80, for FDR corrected t- and U-tests between groups (C, R)). However, the Random group showed a better mean memory consolidation across all force field magnitudes (Posttest ALL: *t*(22) = 3.23, *p* = .016, *d* = 1.32, FDR corrected), with this effect most pronounced predicting high effect sizes using Bayesian statistics (see Supplementary Figure S2).

However, it is important to mention here that this consolidation effect occurred due to performance differences at the end of Practice, for there is no group difference regarding absolute Posttest values (Posttest FT: *U* = 46, *p* = .143, *d* = −0.36; Posttest ALL: *t*(22) = −0.41, *p* = .684, *d* = −0.17, for t- and U-tests between groups (C, R)). This indicates that benefits of the Random group cannot be seen in the absolute Posttest performance. We confirmed this indication by showing that at least two parameters during Practice are having a confounding effect on the absolute Posttest performance across all groups, namely motor error (*r* = 0.56, *p* < .001) and motor variability (*r* = 0.58, *p* < .001, for uncorrected Pearson correlations of all participants (BM, BU, C, R) between Practice LT and Posttest ALL and between the individual’s standard deviation of the whole practice session and Posttest ALL). However, since the Random group revealed a higher motor error (*t*(22) = −2.89, *p* = .009, *d* = −1.18) and a higher motor variability (*t*(22) = −4.22, *p* < .001, *d* = −1.72, for t-tests between groups (R, C)) compared to the Constant group during Practice, correlation coefficients might be compromised due to the inclusion of the Random group (Fig. 3C). A deeper investigation of the data shows that Pearson coefficients of the Random group were indeed higher compared to coefficients of the other groups but differed significantly only for the motor variability (motor error: *z* = −1.28, *p* = .201; motor variability: *z* = −2.63, *p* = .018, for the uncorrected differences between groups (R, [BM BU C]) after r-to-z transformation). This confirms that an increase in motor variability during Practice might increase the motor memory consolidation (from Practice to Posttest) but also confounds the absolute Posttest performance, with this effect being more pronounced in the Random group than in all the other groups.

## 4. Discussion

Our results showed no differences between the two Blocked groups although the Posttest schedule of one group (BM) did and the other schedule (BU) did not control for retroactive inhibition. Compared to the Random group both Blocked groups showed a limited memory consolidation, which depicts that retroactive inhibition does not account for CIE in motor adaptation tasks. Comparisons between Random and Constant groups showed a similar memory consolidation for each single force field magnitude. However, the Random group outperformed the Constant group in its mean memory consolidation across all three force field magnitudes.

### 4.1. Retroactive inhibition does not affect the contextual interference effect in motor adaptation

The experimental procedure of the BM group controlled for possible confounding effects of retroactive inhibition. Nevertheless, BM performed similar to BU and its memory consolidation was hampered compared to the Random group. These findings contradict previous skill learning studies (Shea & Titzer, 1993, Del Rey et al., 1994, Shewokis et al., 1998), assuming retroactive inhibition as the underlying mechanism for the contextual interference effect. Although retroactive inhibition seemed to decrease the Posttest performance in the BU group (Fig. 3A), this effect was too small to explain the benefits after random practice.

These benefits for the Random group were also observed when testing for the generalization of memory to the contralateral hand. This finding concurs with the literature, which frequently showed CIE for transfer tests in skill learning tasks (e.g. Shea & Morgan, 1979, Brady, 2004, Wright et al., 2015) and reproduces earlier findings from our lab using a motor adaptation task (Thürer & Weber et al. 2018). Our results indicate that participants of the Random group were not only able to consolidate better in the meantime between sessions, leading to similar initial performances in the Post- and Transfer test, they also were able to adapt faster towards the force field condition with their left hand. This positive effect of variability on subsequent motor adaptation is in line with a previous study, demonstrating that participants revealing a highly variable baseline period adapt faster during the subsequent practice period (Wu et al., 2014). It is assumed that this positive effect occurred due to noise in the motor planning system but not due to noise in the motor execution system (Dhawale et al., 2017). That leads to the suggestion that the nervous system, at least in some way, uses the knowledge of uncertainty of measured and/or predicted feedback (Izawa & Shadmehr, 2008) to improve motor adaptation (Wei & Körding, 2010). However, future work is needed to investigate this more deeply.

### 4.2. Contextual interference improves only the mean memory consolidation of multiple force field magnitudes

Although CIE reflects a widely accepted phenomenon and seems to be unaffected by retroactive inhibition in motor adaptation tasks, it is not clear whether random practice is always beneficial over constant practice. Our results showed that benefits of random compared to constant practice regarding motor memory consolidation occur only if multiple force field magnitudes are retested. This indicates that memory consolidation of a single task might not be improved by a highly variable practice schedule. This concurs with the especial skill effect for skill learning (Breslin et al. 2010) but contradicts previous work regarding random practice (Shea & Kohl, 1991).

This finding is also in line with our correlation results. We were able to show that both, an increased motor error and an increased motor variability during Practice hamper the absolute Posttest performance. Especially the absolute Posttest performance of the Random group was reduced by the confounding effect of motor variability. However, it is important to note that absolute values of Posttest performance did not differ significantly between groups. Nevertheless, derived from a practical perspective, Random practice might be the better choice of scheduling a practice session since it leads to similar results than constant practice but has the opportunity to increase mean memory consolidation of multiple force field magnitudes and to enhance the generalization, in terms of faster re-adaptation on the contralateral hand (Fig. 3B).

In addition, it might be that a lower amount of motor variability during practice would lead to the same consolidation benefits but would also lead to better absolute performance values of the Random compared to the Blocked or the Constant group. In a previous study, we used a random practice design with lower inter-trial variabilities and were able to show better absolute performance values for Random compared to Blocked groups throughout the whole transfer test (Thürer & Weber et al., 2018). This indicates that the beneficial potential of variable practice depends on the right amount of variability during practice.

### 4.3. Limitations

This study showed some minor limitations, which we would like to address. The Constant group trained the same amount of trials as the other groups but each subject of only one force field magnitude. Therefore, this group was able to draw on a greater practice experience for one specific magnitude compared to the other groups. We cannot state how much this affected the results but from a practical perspective it was important to have the same amount of practice time for each group.

The force field magnitudes might have been too different and, thus, induced a too high practice variability in the Random group. This might be the reason why we were not able to show absolute Posttest and Transfer test performance benefits for the Random group. In a previous study with a lower amount of variability, we were able to show these absolute benefits after Random practice in the transfer test on the contralateral hand (Thürer & Weber et al., 2018).

The order of Post- and Transfer tests was not counter-balanced. Therefore, similar group performances in the first Transfer trials might be caused by the 18 Posttest trials. However, we were previously able to show that contralateral transfer from the dominant to the non-dominant hand after random practice is almost independent of the Posttest performance (Thürer & Weber et al., 2018) and, therefore, suggest that this had only minor effects on our results.

In this study, we investigated motor adaptation and not skill acquisition and, therefore, our interpretations cannot be generalized to skill learning tasks. However, from a theoretical point of view, confounding effects of retroactive inhibition should be more prone to happen in motor adaptation than in skill acquisition, due to a bigger potential overlap of the underlying neural structures.

## 5. Conclusion

In this study, we were able to show that the contextual interference effect represents a valid learning phenomenon that is not affected by retroactive inhibition. Furthermore, we were able to show that benefits of random practice are more related to the memory consolidation of multiple tasks / parameters and to a faster re-adaptation on the contralateral hand. However, variability in general must not always be beneficial regarding a single task / parameter or regarding the absolute performance values in a posttest. However, it remains unsolved how the motor system uses variability to improve subsequent motor memory consolidation, which needs further investigation on the neurobiological level.

## Acknowledgment

We like to thank Alexander Wolpert for his technical assistance and Ernst Hossner for the fruitful discussions and his valuable input.

## Author contribution statement

The study was designed by BT, AF, and TS. SG recorded the data and BT performed the data processing and statistical analysis. BT wrote the first draft of the manuscript and all authors contributed to manuscript revision. All authors read and approved the submitted version of the manuscript.

